# Living in the City: Symbiont stability and bacterial compositional and functional plasticity in Miami’s urban corals

**DOI:** 10.64898/2026.02.25.707149

**Authors:** Thinesh Thangadurai, Anthony J Bellantuono, Daniel G Merselis, R Riley Hatch, Aniruddha Marathe, Colin Foord, Mauricio Rodriguez-Lanetty

## Abstract

Increasing urbanization and climate change pose significant threats to coral reefs, highlighting the need to understand the process underlying coral acclimatization in urban environments. Alterations in the microbiome composition represent a key mechanism by which corals adapt to varying environmental conditions. We compared endosymbionts and bacterial communities associated with *Siderastrea siderea* from urban and offshore Miami reef tracts across three seasons. We found two distinct genera of endosymbiotic Symbiodiniaceae algae, namely *Cladocopium* and *Breviolum,* consistently across sites and seasons, with *Cladocopium* predominating. The stable presence of these symbionts suggests host specificity in *S. siderea* and highlights the potential advantage of harboring multiple symbionts to enhance survival in diverse environments. In contrast, bacterial diversity exhibited variation across seasons and locations, with a small subset of microbes identified as a core microbiome demonstrating the remarkable plasticity of bacterial communities in response to environmental changes. Differential analysis revealed an increased abundance of *Alteromonas* and *Synechococcus* in urban corals, which may contribute to host nutrient acquisition, antibiotic production, and survival in polluted environments. Predicted functional profiles further demonstrated distinct metabolic reorganization of microbial communities between urban and offshore reefs, with urban corals enriched in pathways associated with stress response, pollutant degradation, and nutrient cycling. Together, these findings indicate that while algal symbionts remain stable, bacterial communities undergo both compositional and functional plasticity that likely supports coral persistence in highly urbanized environments.

## Introduction

Coral reefs have declined globally over the past two decades due to anthropogenic climate change and urbanization [1–4]. Urban-driven sedimentation and nutrient enrichment degrade coastal habitats, limiting hard coral survival [5–9]. However, thriving coral populations have been documented in several urban tropical and subtropical regions, including Miami [8,10–13] These coral colonies thrive on artificial substrates such as concrete walls and riprap in the Port of Miami, Star Island, and MacArthur basins. In contrast, corals along the Florida reef tract have declined due to bleaching and disease [14–18]. The persistence of corals in urban habitats highlights their adaptive capacity and underscores the importance of understanding acclimation mechanisms to guide conservation under intensifying urbanization and climate change [19–21].

Coral survival across broad geographic ranges depends on complex microbial consortia, including endosymbiotic dinoflagellates, bacteria, viruses, fungi, and archaea [22–28]. Corals rely heavily on photosynthetically derived metabolites from their endosymbionts [29]. These symbionts exhibit substantial taxonomic and metabolic diversity and, owing to shorter generation times, may enhance host adaptive potential [30, 31]. Community restructuring through “shifting” and “shuffling” can further increase coral plasticity [32–34]. However, the consistency of such restructuring across species remains unclear. For example, *Pocillopora verrucosa* maintains stable Symbiodiniaceae communities across environmental gradients [35], whereas *Porites lutea* exhibits high flexibility [36], indicating species- and genotype-specific responses to environmental change [37].

Beyond Symbiodiniaceae, bacteria are integral to coral health, contributing to nutrient cycling, UV protection, and antimicrobial defense [38–40]. Although bacterial assemblages vary with age and environmental conditions, a conserved core microbiome has persisted for over 425 million years across diverse habitats, including reef pools [41, 33], volcanic craters [42] and deep-sea environments [43]. Reciprocal transplantation studies further demonstrate that shifts in bacterial communities enable acclimatization to novel environments while maintaining this core microbiome [41, 44]. Therefore, characterizing symbiont and bacterial communities in urban corals is essential for understanding microbial mechanisms underpinning resilience in extreme habitats.

Our current understanding of coral responses to environmental stress has largely emphasized interactions with either Symbiodiniaceae or bacteria independently. However, growing evidence indicates that specific bacterial groups contribute to organic mineralization processes that enhance reef resilience to ocean acidification [45, 46]. The co-occurrence of particular bacterial taxa with Symbiodiniaceae further suggests that coral metabolic homeostasis depends on inter-domain interactions [47, 48], and some symbionts may rely directly on bacterial partners for survival [45, 49, 50]. Even low-abundance taxa (2–5%) can exert disproportionate functional influence, underscoring the need to integrate functional analyses with community composition [51–54]. Although most coral microbiome studies document taxonomic shifts across environmental gradients, few assess whether these changes correspond to functional reorganization. Linking community structure to metabolic capacity is therefore essential to elucidate mechanisms of coral acclimatization under anthropogenic stress. Predictive functional profiling offers a powerful framework to evaluate microbial metabolic shifts associated with urbanization, providing mechanistic insight beyond compositional patterns alone.

We hypothesized that (i) corals in urban environments undergo changes in their bacterial and symbiont community composition and associated microbial functions in response to their surroundings, and (ii) these changes vary seasonally. We examined the endosymbiont and bacterial communities associated with coral species in urban and reef tract environments to test these hypotheses. This study was conducted over three seasons—two summers and one winter—at three urban reef sites (less than 1 meter deep as 1 meter was the maximum available depth at these inshore locations) and three reef tract sites (approximately 2.5 - 5 meters deep-the shallowest available). Using culture-independent techniques, we aimed to determine the extent to which bacterial and endosymbiotic communities change to support coral survival in urbanized environments and identify specific bacterial community functional reorganisation/changes necessary for coral survival in these settings. Additionally, we investigated seasonal variations in the endosymbiont and bacterial communities, as well as specific bacterial species in *Siderastrea* corals. Lastly, we sought to identify the core microbiome of *Siderastrea* corals, examining its consistency across seasons and locations, regardless of the urbanized environment.

## Methodology

### Sampling sites

Sampling was conducted across six sites during three seasons (August 2017, September 2017, and January 2018). These included three urban inshore sites—North MacArthur (NM), South MacArthur (SM), and Star Island (STR)—where corals occurred on rocks near the bus stand, and three offshore Florida Reef Tract sites—Miami Beach (MB), Emerald Reef (EM), and Second Reef (SR)—located 5.5–8 km offshore (∼4 km from the coastline; Fig. 1).

**Fig. 1.**
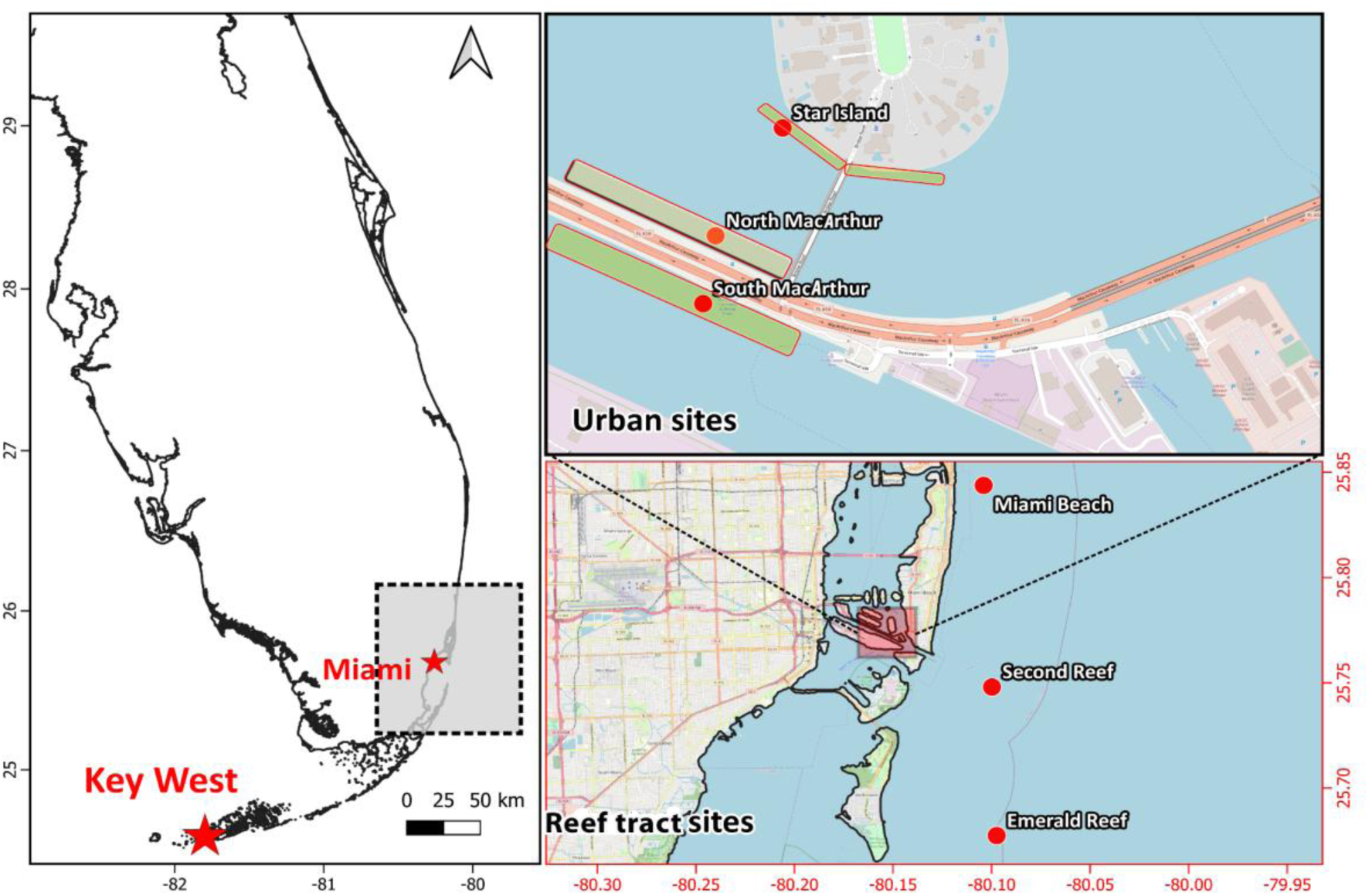
Map of study sites in Reef tract, Selected three urban sites (Top right). Selected reef tract study sites (Bottom Right).

Urban sites were directly influenced by anthropogenic activities. Watershed nutrient data obtained from the Florida Department of Environmental Protection (2025) indicated increasing inorganic nitrogen and phosphorus concentrations during the eight years preceding sampling. Concurrently, long-term monitoring records showed declining chlorophyll-a concentrations across reef sites [55, 56, 57]. Coral communities were observed thriving in anthropogenically modified coastal environments along the roadside near Miami Beach (Fig. 2a). A well-developed colony of *Colpophyllia natans* was documented within this artificial setting, exhibiting typical massive growth morphology (Fig. 2b). Additionally, colonies of *Siderastrea siderea* were recorded at Star Island, where they displayed healthy surface texture and structural integrity despite the modified coastal conditions (Fig. 2c&d).

**Fig 2.**
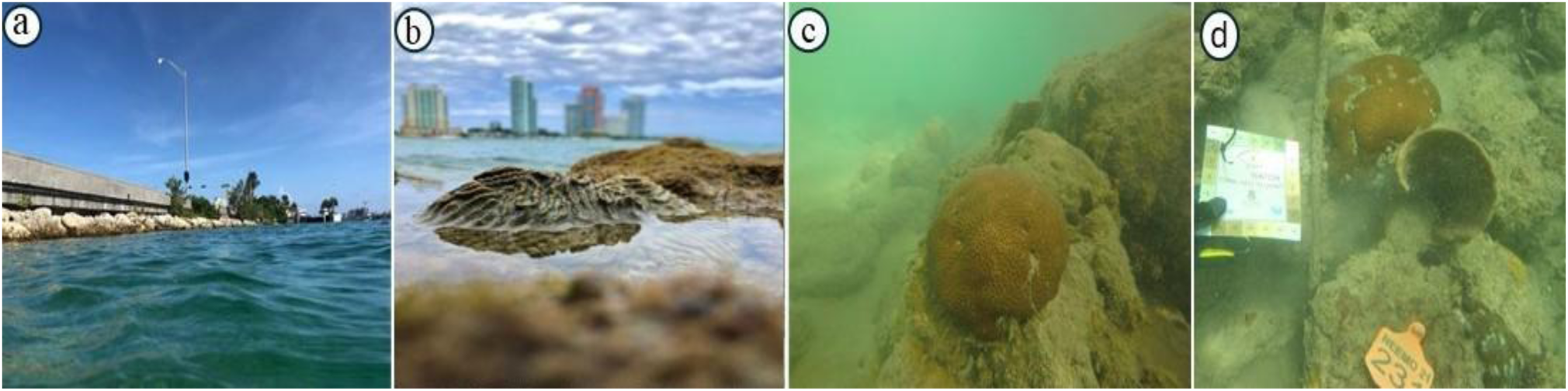
a) Corals inhabiting man-made habitat along the road near Miami Beach. b) *Colphophylia* colony in the manmade habitat (c & d) *Siderastrea siderea* colony at Star Island

### Sample collection

*Siderastrea siderea* was selected because it occurred at all sites. A total of 180 samples were collected (10 per site per season) from urban (0.3–1 m depth) and reef tract sites (2–3 m depth; shallowest accessible). This design was repeated across three seasons for symbiont analyses. For bacterial 16S rRNA gene profiling, six samples per site were selected (36 per season), totaling 108 samples across seasons. Coral tissues were collected, preserved in a DMSO buffer within 2 mL vials, stored at 4°C, and subsequently transferred to −20°C until DNA extraction. Endosymbiont and bacterial analyses followed Rodriguez-Lanetty et al. (2013) [58].

### DNA extraction, amplification, and sequencing

DNA was extracted using the MP FastDNA Spin Kit for Soil (MP Biomedicals, USA) with an additional ethanol precipitation step. Extracted DNA was divided into two aliquots for bacterial 16S rRNA and Symbiodiniaceae cp23S rRNA sequencing. The ∼480 bp cp23S region was amplified using primers F:AATAACGACCTGCATGAAAC and R: GCCTGTTATCCGTAGAGTAGC with GoTaq polymerase. Although GoTaq has a higher error rate than high-fidelity enzymes, it was selected because alternative polymerases failed to produce consistent, specific amplification despite optimization of temperature, MgCl₂, and template concentrations; potential errors were addressed during downstream analysis.

PCR reactions (20 μL) contained 10 μL 2× GoTaq Master Mix, 1 μL DNA template, 0.5 μL each forward and reverse primers, 1.2 μL 25 mM MgCl₂, and 6.8 μL molecular-grade water. Thermal cycling consisted of 95 °C for 3 min; 35 cycles of 95 °C for 30 s, 55 °C for 30 s, and 72 °C for 30 s; followed by a final extension at 72 °C for 5 min. PCR products were purified using magnetic beads according to the Illumina 16S amplicon protocol. Amplicons were indexed using KAPA HiFi HotStart ReadyMix with an initial 95 °C for 3 min, followed by 10 cycles of 95 °C for 30 s, 55 °C for 30 s, and 72 °C for 30 s, and a final extension at 72 °C for 5 min. Libraries were purified, normalized using SequalPrep plates, and sequenced (300 bp paired-end) on an Illumina MiSeq at Florida International University. Additional library preparation indexing was performed using the HOTStarTaq Plus Master Mix kit (Qiagen). The cp23S PCR program consisted of 95 °C for 3 min; 25 cycles of 95 °C for 30 s, 55 °C for 30 s, and 72 °C for 30 s; and a final extension at 72 °C for 5 min.

The bacterial 16S rRNA gene (V3–V4 regions; ∼550 bp) was amplified using primers containing Illumina overhang adapters: Forward (5′ TCGTCGGCAGCGTCAGATGTGTATAAGAGACAG CCTACGGGNGGCWGCAG) and Reverse (5′ GTCTCGTGGGCTCGGAGATGTGTATAAGAGACAG GACTACHVGGGTATCTAATCC), synthesized by Integrated DNA Technologies (USA). PCR reactions (25 μL) contained 12.5 μL KAPA HiFi HotStart ReadyMix, 5 μL of each primer (1 μM), and 2.5 μL template DNA (5 ng/μL). Cycling conditions were 95 °C for 3 min; 30 cycles of 95 °C for 30 s, 55 °C for 30 s, and 72 °C for 30 s; followed by 72 °C for 5 min. Amplicons were purified using AMPure XP beads. For indexing, 5 μL purified amplicon was subjected to 8 PCR cycles with dual P5/P7 indices, followed by bead cleanup. Libraries were quantified using a Qubit 3 fluorometer (dsDNA HS Assay), pooled equimolarly, and quality-checked by 1% agarose gel and Agilent 2100 Bioanalyzer prior to sequencing with MiSeq v3 (600-cycle) reagents.

### Sequence processing and selection of operational taxonomic units

Paired-end reads were quality- and primer-trimmed in QIIME 2 [59]. Overlapping reads were merged and denoised using the DADA2 workflow to generate ASVs [60]. Taxonomic assignment was performed with the QIIME2-based RDP classifier against the SILVA database [61]. Sequences unclassified at the kingdom level and non-bacterial reads (mitochondria, chloroplasts, archaea, and eukarya) were removed. ASVs with <20 total reads were excluded to generate the final ASV table used for downstream analyses. Alpha diversity metrics (Chao1, Shannon diversity, Simpson’s evenness, and phylogenetic diversity) were calculated in QIIME 2.

### Assessing microbial biodiversity & core microbiome

Paired-end reads were trimmed in QIIME 2 and denoised with DADA2 to generate 16S rRNA ASVs [59]. Taxonomy was assigned using the RDP classifier against SILVA [61, 62]. Contaminants, unclassified sequences, and non-bacterial reads were removed. Alpha diversity (Faith’s PD, Pielou’s evenness) and beta diversity (weighted and unweighted UniFrac) were calculated in q2-diversity [59, 63–65]. PCoA was performed in R (‘Vegan’) [66], and boxplots were generated in MicrobiomeAnalyst [67]. The core microbiome was defined as ASVs present in ≥95% of samples. Analyses were conducted in R v4.2.1 [68].

### Predicted functional profiling of microbial communities

Predicted functional profiles of coral-associated bacterial communities were inferred from 16S rRNA gene sequencing data using a bioinformatics-based functional prediction approach (PICRUSt2 pipeline implemented within MicrobiomeAnalyst v2.0). Taxonomic profiles were mapped to Kyoto Encyclopedia of Genes and Genomes (KEGG) Orthology groups to estimate the relative abundance of metabolic pathways across samples. Functional predictions were normalized to account for variation in 16S rRNA gene copy number prior to pathway reconstruction. Predicted pathway abundances were subsequently compared between urban and offshore reef sites and across seasons to assess functional reorganization associated with urbanization. Inferred functional profiles were analyzed using NMDS ordination and PERMANOVA to evaluate differences across locations, sites, and sampling seasons.

### Symbiodiniaceae 23s amplicon sequencing library preparation

Symbiodiniaceae amplicon libraries targeting the chloroplast 23S rDNA hypervariable region (∼480 bp) were prepared and sequenced (300 bp paired-end) on an Illumina MiSeq at the FIU Forensics Sequencing Facility. Amplification used universal primers F: AATAACGACCTGCATGAAAC and R: GCCTGTTATCCGTAGAGTAGC. Reads were demultiplexed with Cutadapt and processed in QIIME 2 for quality and primer trimming. Sequences were denoised using DADA2 to generate ASVs and remove chimeras. Taxonomic classification was performed using a custom classifier constructed with the QIIME RESCRIPt pipeline based on publicly available 23S Symbiodiniaceae sequences retrieved from NCBI (May 2023) [69]. Sequences unclassified at the genus level and ASVs with <20 total reads were removed prior to downstream analyses.

## Result

### *Symbiodiniaceae* community structure

High-quality reads were obtained from 154 samples, yielding 1,038,355 sequences prior to filtering (mean: 6,742 reads per sample). High-quality reads were defined as sequences retained after DADA2 processing in QIIME2, including quality trimming, chimera removal, and denoising. Two dominant symbiont genera, *Cladocopium* (87%) and *Breviolum* (13%; predominantly *B. minutum*), were detected across samples (Fig. 3). Minor taxa included *Breviolum psychrophilum* and an unclassified *Cladocopium*, each comprising <0.05% of reads. Overall, 63% of *Siderastrea* samples hosted only *Cladocopium*, whereas 37% contained both genera; among mixed colonies, 88% were *Cladocopium*-dominated and 12% *Breviolum*-dominated. Endosymbiont community structure remained stable across sites, locations, and seasons, with no significant differences detected by PERMANOVA (F = 1.9, R² = 0.011, p = 0.059).

**Fig. 3.**
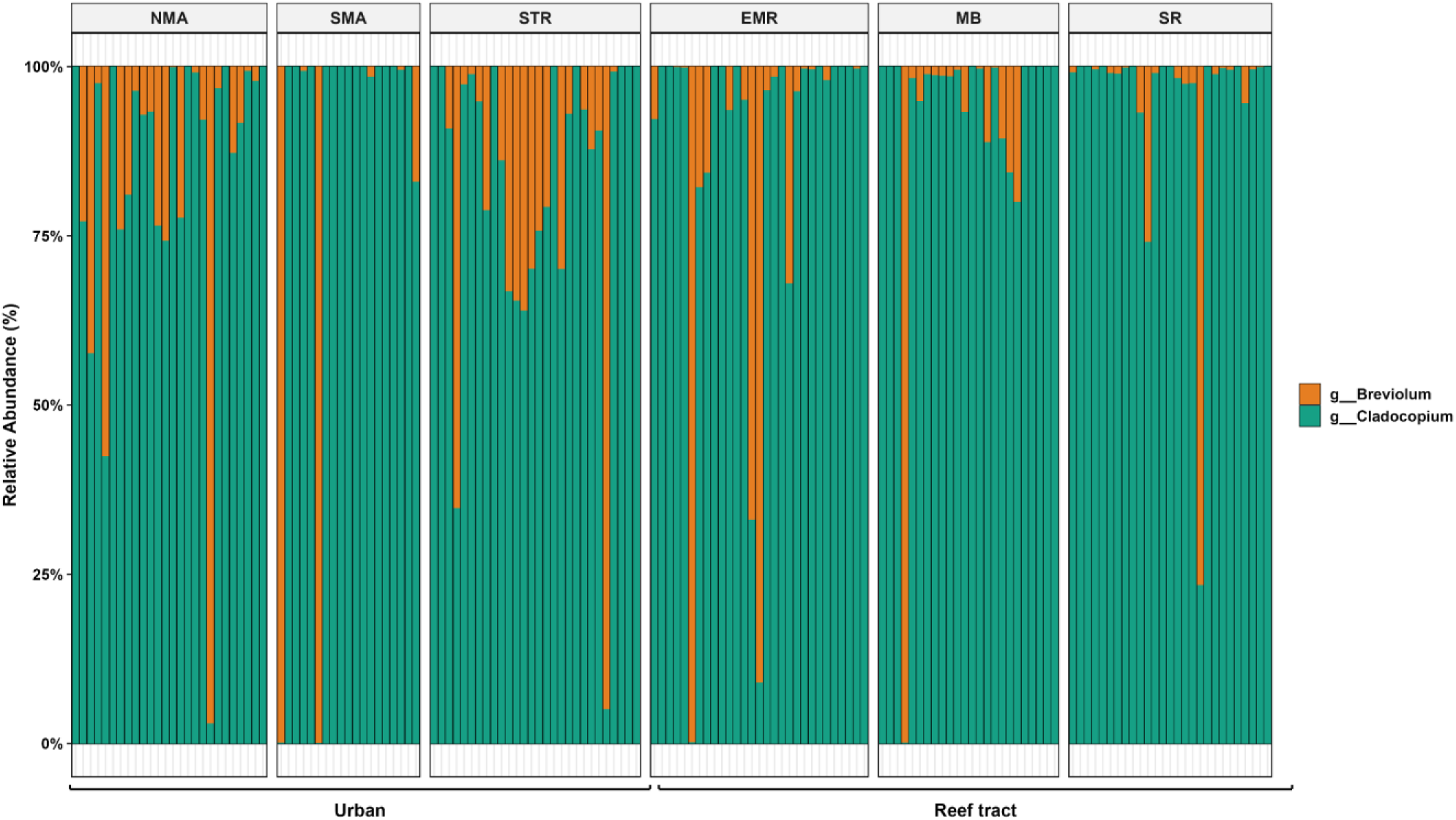
Taxonomic profile (genus level) of abundant symbiont communities associated with *Siderastrea* samples collected from six surveyed sites in Miami’s urban (left) and offshore reef tract (right) environments. *Cladocopium* and *Breviolum* were the dominant symbiont genera observed across both site types.

### Bacterial Community Structure

Bacterial 16S rRNA gene amplicon sequencing from *Siderastrea* samples from urban and offshore locations (*n* = 97) yielded 7,377,784 sequences (median = 55286 reads). Sequence length ranged from 260 to 447 bp (median = 398 bp). 4,257 ASVs were recorded across all samples, ranging from 159 to 2,556 ASVs per sample (median = 656). Out of 4,257 ASVs, The ten most dominant ASVs comprised 42% of the total bacterial community abundance. The remaining ASVs were rare. (i.e., each contributed < 1% of total abundance), but shaped the remaining microbial community structure (39.1%) without defined dominant taxa (Fig. 4). The most abundant bacterial phylotypes at the genus levels were a single *Ralstonia* sp. (14.0%), Vibrio (7%) *Alteromonas* (7%), *Pseudoalteromonas* (7%) and *Erythrobactor* 2.8% (Fig. 2). In family level Burkholderiaceae (11%), Rhodobacteraceae (7.8%), Vibrionaceae (7%), and Bacteriodota (6.8%) and Alteromonadaceae (6.1%). Finally, unclassified ASVs at the phylum level, and those with less than 20 reads across the entire dataset yielded 4257 final ASVs assigned to 568 different taxa at the genus level.

**Fig 4.**
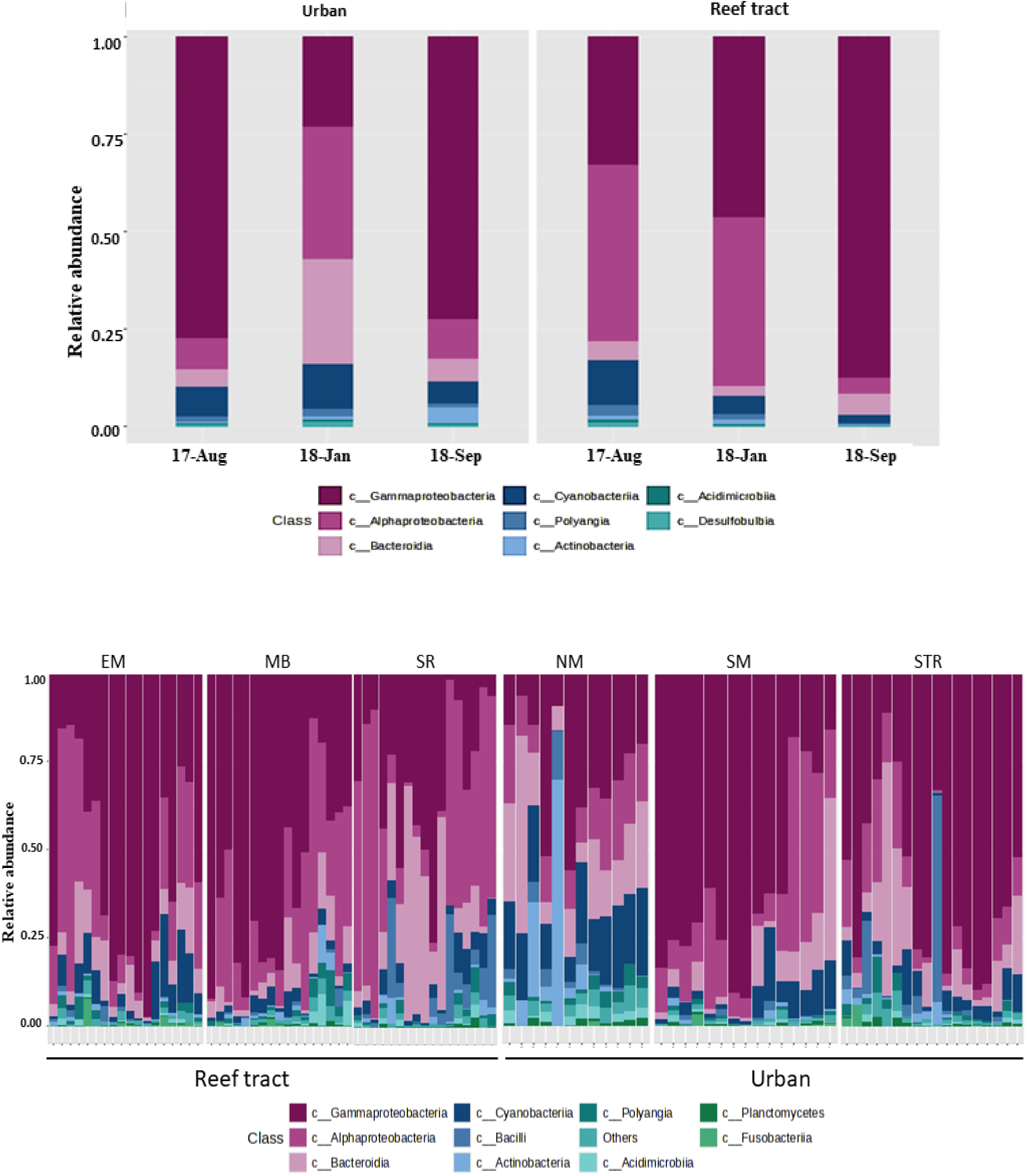
Abundant bacterial community taxonomic profile (genus level) associated with *Siderastrea* samples collected from 6 surveyed sites in the Miami urban sites (Left) and reef tract (Right). *Ralstonia* and *Vibrio* were the most dominant ASVs and composed 43.6% of the total community in both sites. The bacterial community was significantly different between seasons, not between sites.

### Bacterial Community Analysis

Alpha diversity (Faith’s PD) did not differ significantly between urban and offshore reef tract locations (Kruskal–Wallis, p > 0.05). However, pairwise comparisons showed significant differences across seasons (p < 0.0002) and sites (p < 0.003). Beta diversity analyses using weighted and unweighted UniFrac revealed significant compositional differences between urban and offshore locations (unweighted: F = 2.0, p < 0.05) and significant effects of location (F = 6.5, p < 0.05), season (F = 2.0, p < 0.05), and site (F = 2.0, p < 0.05) under weighted UniFrac (PERMANOVA). PCoA based on Bray–Curtis dissimilarity indicated that seasonal variation primarily explained bacterial community differences, with no significant separation among sites (Fig. 5).

**Fig 5:**
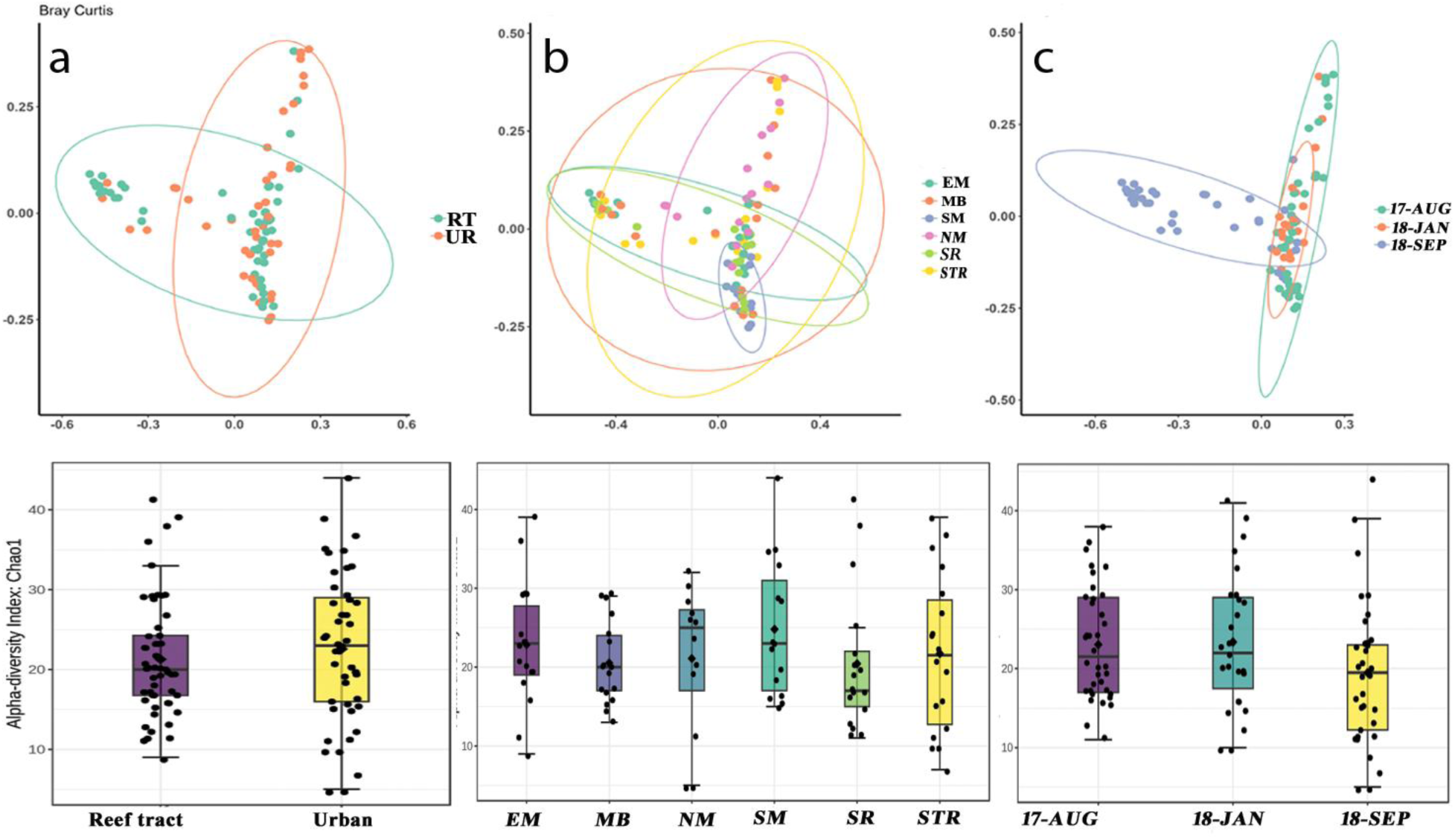
Shannon index value for location, Sites, and Season. Principal coordinate analysis (PCoA) based on the Bray-Curtis dissimilarity matrix of bacterial communities associated with *Siderastrea siderea* from six sites (three Urban sites, three offshore reef tract sites, and three different seasons). PCoA shows a clustering pattern (a) between different Locations (b). between sites, c) between seasons. Compositional differences in bacterial communities were best explained by seasons.

### Differential analysis

Differentially abundant taxa between urban and offshore reef tract samples were identified using LEfSe based on 16S rRNA taxonomic profiles. LDA scores and p-values were calculated for each taxon. Ten ASVs differed significantly between groups (Table 1). Sphingomonadaceae was significantly enriched in offshore samples (LDA = 4.5, p = 0.001), whereas Alteromonadaceae was enriched in urban corals (LDA = 4.7, p = 0.001). These taxa were among the most discriminative features between habitats. These findings highlight distinct microbial signatures associated with urban and offshore habitats and demonstrate the utility of LEfSe in identifying taxa discriminating between environmental conditions (Fig. 6).

**Fig 6:**
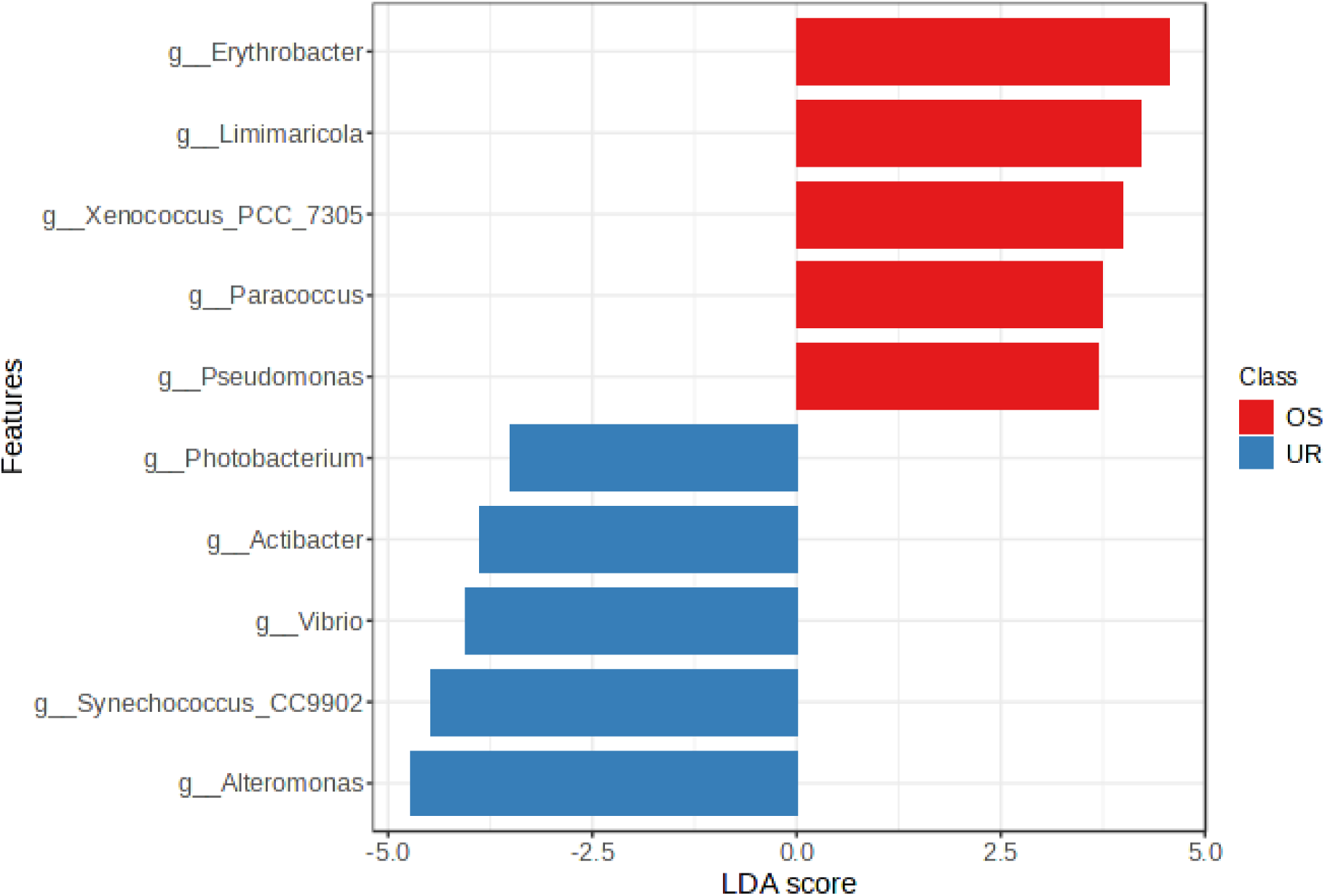
Observed taxonomic discrimination in microbiome composition between urban and reef tract corals revealed by LEfSe analysis.

### The core microbiome

Despite high bacterial diversity across samples (4,257 phylotypes), only a small subset formed a consistent core microbiome across sites. Six phylotypes were detected in ≥95% of coral individuals (core microbiome; Fig. 4), while 175 occurred in ≥50% of colonies, indicating that most taxa were spatially variable.

Core taxa were dominated by *Ralstonia* (61.9%), followed by *Pseudomonas* (17.5%) and *Pseudoalteromonas* (10.3%), with minor contributions from Cyanobacteria (1%) and Firmicutes (2.1%). Within Proteobacteria, Gammaproteobacteria comprised 53.3% of core OTUs, whereas Alphaproteobacteria (16.7%), Deltaproteobacteria (13.3%), Epsilonproteobacteria (8.3%), and Betaproteobacteria (6.7%) were less represented.

Among the six highly persistent phylotypes (Fig. 5), four were assigned to *Ralstonia* and *Gluconacetobacter*, while the remaining taxa were classified at higher taxonomic levels. Notably, the most prevalent phylotypes, OTUs 306 and 25296, belonged to Bacteroidetes and Deltaproteobacteria (Fig 4), respectively, and represent taxa infrequently reported in coral-associated microbiomes (Fig 7).

**Fig 7:**
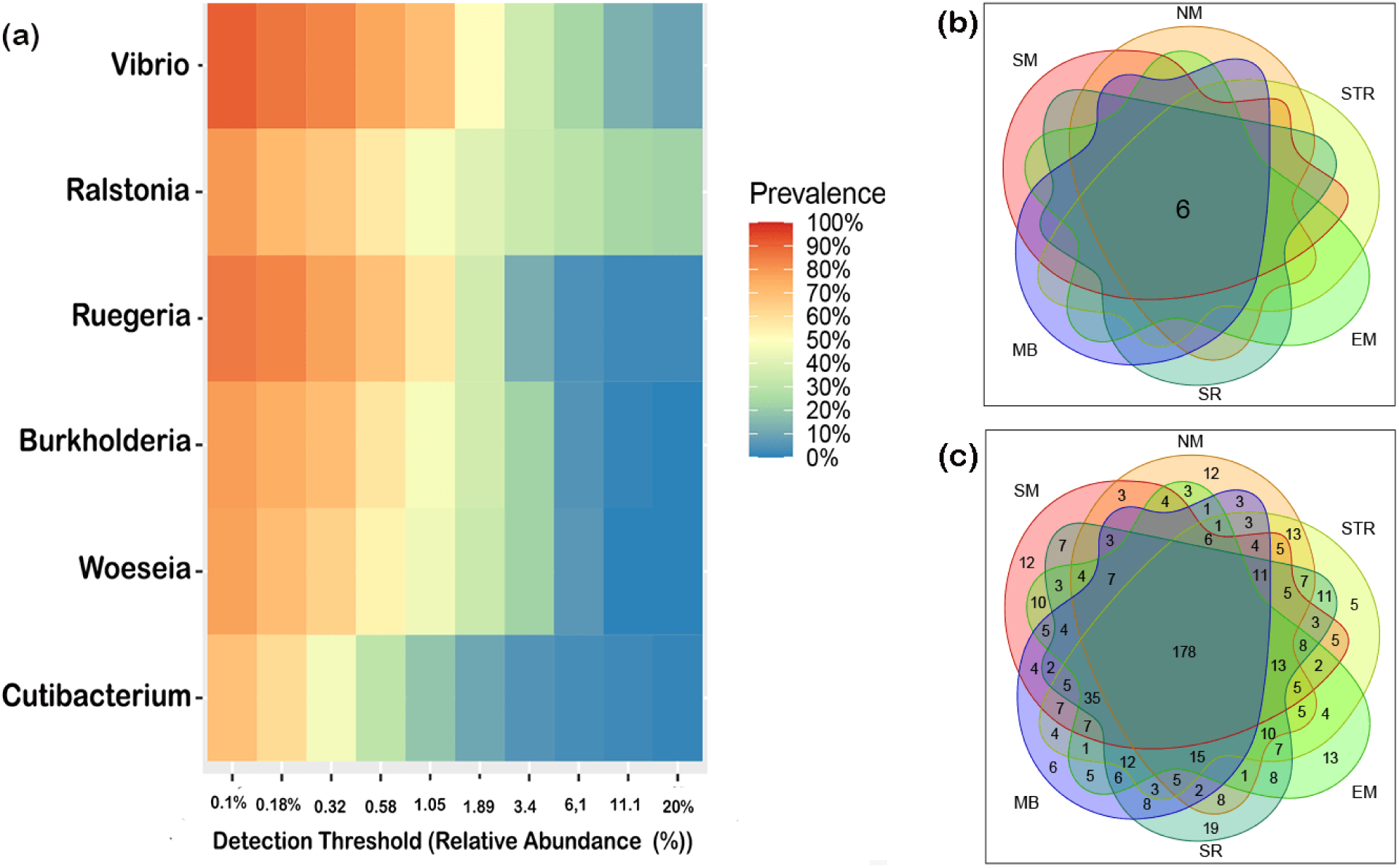
Heat Map:- (a)The x-axis represents the detection thresholds indicated. The heat map of the bacterial genus core. The x-axis represents the detection thresholds (indicated as relative abundance) from lower (left) to higher (Right) abundance values. Color shading indicates the prevalence of each bacterial family among samples for each abundance threshold. As we increase the detection threshold, the prevalence decreases. Venn diagram showing the number of shared bacterial genera across sites without any filter. (b) shows six genera in over 95% of coral individuals from each site. (c)178 genera in over 50% of corals.

### Variation in Specific Predicted Gene Functions Across Sampling Locations

Predicted functional pathway profiles differed significantly between urban and offshore reefs (Fig. 8; Supplementary Table 1). Pathways were considered significantly associated with location at p < 0.05, indicating that observed differences were unlikely due to random variation. This analysis aimed to evaluate how environmental conditions and anthropogenic inputs influenced predicted microbial functional profiles.

**Fig 8:**
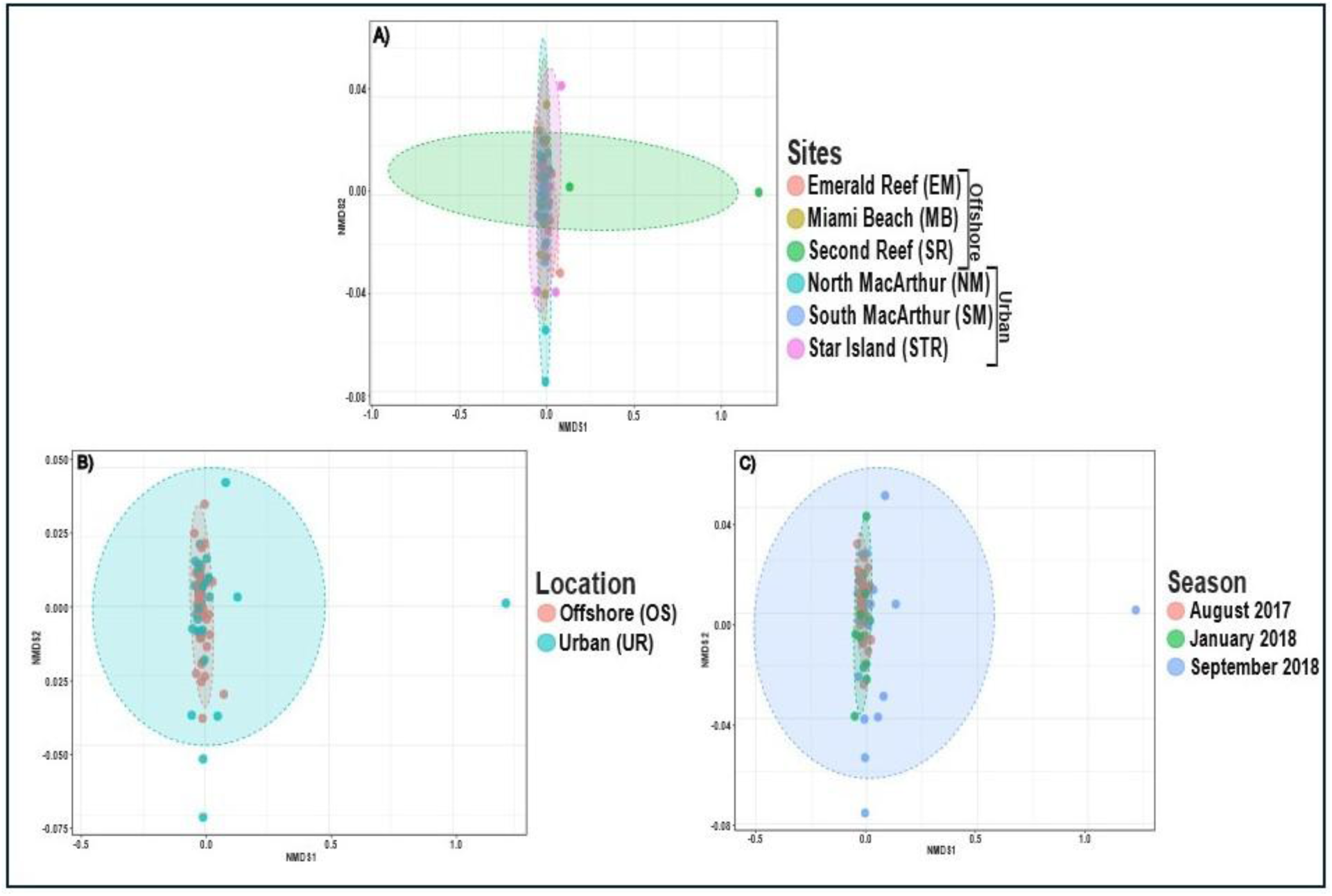
Inferred functional profile NMDS plots for *S. siderea* samples by A) reef site (oval colors correspond to communities at each individual site, with offshore/urban locations encompassing sampling sites indicated as vertical labels in legend), B) location (ovals correspond to offshore/urban),, and C) sampling season/date (ovals correspond to individual collection efforts). NMDS stress value=0.091214, quantitative PERMANOVA results indicating significance of difference between factors are found in Table 3.4, below. Plot images generated in <u>microbioeanalyst.ca</u> v. 2.0 and edited in Inkscape v. 1.4.2.19

Both reef types exhibited 134–136 pathways across ∼2,400 database hits (±4%). In both environments, genes associated with photosynthesis were significantly represented (p = 0.010–0.02), along with pathways related to amino acid, carbohydrate, and methane metabolism.

Urban reefs showed greater enrichment of pathways linked to toxin processing, waste mitigation, and stress-related metabolism compared to offshore reefs. Significantly enriched pathways included methane metabolism, steroid hormone biosynthesis, xenobiotic/drug metabolism, and aminobenzoate degradation. These pathways are commonly associated with pollutants such as fossil fuel derivatives and sunscreen-related contaminants. The complete list of significant pathways is provided in Table 3.2. Collectively, these functional signatures suggest that urban corals persist under environmentally suboptimal conditions and exhibit microbial stress-response mechanisms [70, 71, 72].

PERMANOVA analysis of inferred functional profiles (Supplementary Table 1) showed significant effects of location, site, and season, with only a few non-significant site comparisons (Table 3.4). Non-significant differences were observed among SM and STR (urban), EM, SR, and MB (offshore), and between SM–MB and STR–MB. NMDS ordination (Fig. 3.8A) further indicated greater functional variability within urban reefs compared to offshore sites, explaining the observed dispersion patterns.

## Discussion

The flexibility of coral microbiomes enables corals to tolerate and adjust to environmental stressors, including thermal stress and urbanization [34, 73, 74]. Investigating coral–microbe interactions in urban environments therefore provides critical insight into mechanisms underlying coral resilience. In this study, we examined the roles of endosymbionts and bacterial communities in supporting the persistence of *Siderastrea siderea* within the Miami urban reef system, a habitat exposed to eutrophication, dredging, sedimentation, thermal variability, and high turbidity [75, 76, 77, 78]. Our results identify distinct features of microbiome composition and structure that likely contribute to the ability of *S. siderea* to withstand chronic anthropogenic stress in urbanized coastal environments.

### Stable Endosymbiotic Partnerships Underlying Coral Resilience in Urban and Offshore Habitats

Symbiodiniaceae distribution patterns are shaped by environmental conditions and seasonal variation across broad spatial scales [79, 80, 35]. Corals can respond to local stress by switching or shuffling symbionts to optimize physiological performance under changing conditions [81–83]. Seasonal fluctuations in symbiont abundance and dominance have been widely documented [84–86]. For example, *Orbicella annularis* exhibits biogeographic partitioning of symbiont communities along a ∼1,800 km Caribbean gradient, primarily driven by historical thermal regimes rather than host genetics [87]. Similarly, *Durusdinium* often dominates corals in thermally extreme or marginal habitats, whereas *Cladocopium* is typically associated with cooler fore-reef environments [88, 89]. In addition to environmental factors, host species identity strongly structures symbiont communities across bioregions [90–93].

In contrast to previous studies, our results show that neither site (urban vs. offshore) nor season significantly influenced algal symbiont diversity or composition in *Siderastrea siderea*. This indicates that shifts in Symbiodiniaceae communities are unlikely to be a primary mechanism underlying this species’ persistence in urban environments. Most colonies exhibited mixed symbiosis, dominated by *Cladocopium* with *Breviolum* as a secondary symbiont, while a minority hosted only *Cladocopium*. This pattern is consistent with earlier reports for *S. siderea* and other coral taxa across environmental gradients [83, 94–97]. The recurrent association of *Cladocopium* and *Breviolum* supports their role as generalist Caribbean symbionts [98]. Together, these findings suggest that endosymbiont composition in *S. siderea* remains stable across environments and does not confer substantial adaptive flexibility under urban stress.

The coexistence of multiple symbiont types may nonetheless enhance functional redundancy, buffering corals against environmental disturbances [83, 99–101]. When one symbiont type is compromised by stress or disease, others can maintain key physiological processes such as nutrient exchange and energy production [83, 102]. This redundancy may represent a key mechanism of ecological resilience, allowing *S. siderea* to persist in diverse habitats, from offshore reefs to nearshore mangroves and urbanized coastlines. Thus, the consistent symbiont assemblage observed across the Miami urban and reef tract sites implies that coral resilience in these environments may depend more on host physiological traits or other ecological factors, rather than on symbiont-driven adaptation.

The prevalence of *Cladocopium* in *S. siderea* observed here mirrors patterns documented across the Caribban, including in the Bahamas, Belize, Curaçao, and St. Croix [82, 92, 103]. Thornhill et al. (2006) similarly found stable *Cladocopium*-dominated associations across spatial and seasonal gradients in *S. sidereal* throughout the Florida Keys, highlighting strong host-specific selection for this genus [104]. The dominance of *Cladocopium* in both urban and offshore *Siderastrea* populations underscores the persistence of this symbiosis and may reflect a connection to the broader Florida reef system, where Clade C predominates and is associated with enhanced tolerance to thermal and anthropogenic stress [105]. Collectively, these results underscore the stability of *Siderastrea*-*Cladocopium* partnerships and their potential contribution to coral persistence across increasingly human-impacted marine environments.

### Environmental Variability Shapes Coral-Associated Bacterial Communities Across Reef Habitats

In contrast to the stability of endosymbiont communities, bacterial beta diversity varied significantly between urban and offshore reefs and across seasons. These patterns are consistent with previous studies showing that bacterial communities dynamically restructure in response to environmental drivers such as temperature, salinity, pH, nutrient enrichment, and eutrophication [106–111]. The observed compositional differences highlight strong environmental regulation of bacterial assemblages rather than strict host specificity. Such restructuring likely provides functional advantages to the coral holobiont, facilitating adaptation to local conditions [112–114]. Coral-associated bacteria perform essential roles, including nitrogen fixation, sulfur cycling, and pathogen defense [115–117]. Accordingly, microbiome restructuring is increasingly recognized as a key mechanism underpinning coral plasticity and resilience. Differential abundance analyses further revealed distinct bacterial signatures between urban and offshore reefs, suggesting selective enrichment of taxa that may enhance coral survival under contrasting environmental and thermal regimes [114].

### Urbanization Drives Divergent Bacterial Communities but Retains a Stable Core Microbiome in *Siderastrea siderea*

Differential abundance analyses revealed clear site-specific patterns in the bacterial communities associated with *Siderastrea* siderea. Alteromonadaceae and Photobacterium were enriched in urban reefs, whereas Sphingomonadaceae, *Exiguobacterium, Endozoicomonas, Zunongwangia, Paracoccus, Erythrobacter, and Limimaricola* were more prevalent offshore. These patterns indicate strong environmental filtering, with local conditions likely selecting bacterial taxa that support coral persistence in urban habitats. Despite these compositional differences, a stable core microbiome was consistently detected across all samples. Core taxa included *Ralstonia, Vibrio, Ruegeria, Burkholderia, Cutibacterium, and Woeseia*, suggesting conserved roles in host physiology irrespective of habitat. Although several of these genera are recurrently reported as coral core members, their functional contributions remain incompletely understood. For example, Ralstonia has been proposed to interact with zooxanthellae, potentially facilitating ion-coupled transport and amino acid metabolism within the symbiosis [118], yet direct functional evidence is limited. Given their widespread occurrence across contrasting environments, further investigation is essential to clarify the role of these core taxa in coral acclimatization and long-term adaptation.

### Functional Shifts in Coral-Associated Bacteria Reflect Ecological Adaptation to Urbanization

*Siderastrea siderea* hosts diverse microbial phyla that contribute to key metabolic processes, including nitrate reduction, aromatic compound degradation, and sulfur, ammonia, and carbon oxidation [119–121]. In offshore reefs, *Erythrobacter* emerged as a prominent indicator taxon. This genus contains bacteriochlorophyll-a and carotenoids [122–124], pigments known to regulate reactive oxygen species and support coral physiological stability [125, 126]. Such associations likely reflect stable microbial partnerships under relatively undisturbed reef conditions.

At the genus level, *Ralstonia* and *Vibrio* were among the dominant phylotypes. *Ralstonia* has been reported across multiple coral genera and regions—including the Caribbean, Red Sea, Taiwan, and Hawaii—with relative abundances ranging from 0.5–24%, supporting a potential symbiotic role within the coral holobiont [127–130]. *Vibrio* species are widely detected in coral microbiomes and exhibit dual ecological roles, functioning as opportunistic pathogens as well as beneficial symbionts involved in nitrogen cycling and host metabolism [131–133]. Their recurrent detection across coral taxa and regions suggests a natural association with healthy microbiomes, though their functional versatility warrants further investigation [131].

In contrast to offshore reefs, urban corals exhibited increased relative abundance of *Alteromonas*, *Photobacterium*, and *Synechococcus*. The enrichment of *Alteromonas* suggests adaptive microbial restructuring under urban stress. This genus contributes to DMSP metabolism, antimicrobial compound production, oxidative stress mitigation, and in some strains, nitrogen fixation—potentially supplying fixed nitrogen to algal symbionts and enhancing resilience under nutrient limitation [115, 131, 134–136]. *Photobacterium* supports coral health through antibiotic production, nitrogen fixation, and chitin degradation, conferring both defensive and competitive advantages to the holobiont [137–139]. Similarly, *Synechococcus* contributes to nitrogen fixation and photosynthesis, reinforcing internal nutrient cycling [140, 141]. Its preferential association with urban corals may represent an adaptive strategy to secure alternative nutrient sources under anthropogenic stress [142]. Collectively, these shifts in bacterial composition likely reflect environmentally driven selection of functionally beneficial taxa, providing metabolic and protective advantages that support coral persistence in urbanized habitats [138].

### Inferred Functional Profiles

Taxonomic differences between urban and offshore reefs were accompanied by pronounced shifts in inferred functional capacity. KEGG Orthology analyses revealed distinct functional profiles between habitats. Offshore reefs were enriched in taxa associated with balanced metabolic processes, including Cyanobacteria and *Erythrobacter*, which contribute to nitrogen cycling and biogenic substrate production [143, 144]. In contrast, urban reefs showed enrichment of taxa linked to nutrient-rich and stress-prone environments, including *Synechococcus*. Temporal patterns indicated early enrichment of opportunistic Cytophagales, followed by increased dominance of *Ralstonia* and the unexpected detection of *Pseudomonas yamanorum*, a psychrotolerant species rarely reported in tropical systems. Its presence may reflect previously unrecognized functional plasticity or strong environmental filtering. Although functional pathways were predicted rather than directly measured, these patterns suggest microbiome functional reorganization under anthropogenic disturbance, potentially facilitating coral persistence under suboptimal conditions.

## Conclusion

This study significantly enhances our understanding of the microbiome communities associated with *Siderastrea siderea* corals in urban environments. Our findings show that the Symbiodiniaceae community, particularly *Cladocopium* and *Breviolum*, remains consistent across urban and offshore reefs, suggesting that symbiont composition is not the primary driver of acclimatization between these environments. However, symbiont redundancy might help corals to maintain resilience across diverse environmental conditions. In contrast, we observed significant variations in bacterial community structure between urban and offshore sites and across seasons, indicating that bacterial communities play a more dynamic role in coral adaptation. Urban corals showed an increased abundance of bacterial taxa such as *Alteromonas* and *Synechococcus*, which likely contribute to coral resilience by providing critical functions like nitrogen fixation, disease resistance, and protection against environmental stressors. Consistent with these taxonomic shifts, inferred functional profiles revealed metabolic reorganization of microbial communities in urban reefs, with enrichment of pathways associated with stress response, pollutant processing, and altered nutrient metabolism. These findings highlight the potential role of bacterial flexibility in supporting coral survival in urban environments. Future research should explore the functional roles of these bacterial communities to better understand how they contribute to coral resilience under increasing environmental pressures, including urbanization and climate change.

## Data Availability Statement

The 16S rRNA gene amplicon sequencing data supporting this study have been deposited in the NCBI Sequence Read Archive (SRA) under BioProject accession number PRJNA1424437.

## Acknowledgements

The authors gratefully acknowledge tFlorida Fish and Wildlife Conservation Commission Division of Marine Fisheries Management 620 S. Meridian St., Mail Station 4B3, Tallahassee, Florida 32399-1600, for granting permissions to conduct field sampling and collect coral samples (Licence No - SAL 17 1916-SRP). We thank Florida International University for logistical, laboratory, and sequencing support that enabled this work. TT gratefully acknowledges Sultan Qaboos University for providing research facilities and institutional support. Computational analyses were supported by the HPC Luban Centre at Sultan Qaboos University.

